# Toxicants Associated with Spontaneous Abortion in the Comparative Toxicogenomics Database (CTD)

**DOI:** 10.1101/755868

**Authors:** Sean M. Harris, Yuan Jin, Rita Loch-Caruso, Ingrid Y. Padilla, John Meeker, Kelly M. Bakulski

## Abstract

**Background:** Up to 70% of all pregnancies result in either implantation failure or spontaneous abortion (SA). Many events occur before women are aware of their pregnancy and we lack a comprehensive understanding of high-risk SA chemicals. In epidemiologic research, failure to account for a toxicant’s impact on SA can also bias toxicant-birth outcome associations. Our goal was to identify chemicals with a high number of interactions with SA genes, based on known toxicogenomic responses.

**Methods:** We used reference SA (MeSH: D000022) and chemical gene lists from the Comparative Toxicogenomics Database in three species (human, mouse, and rat). We prioritized chemicals (n=25) found in maternal blood/urine samples or in groundwater, tap water, or Superfund sites. For chemical-disease gene sets of sufficient size (n=13 chemicals, n=20 comparisons), chi-squared enrichment tests and proportional reporting ratios (PRR) were calculated. We then cross-validated enrichment results. Finally, among the SA genes, we assessed enrichment for gene ontology biological processes and for chemicals associated with SA in humans, we visualized specific gene-chemical interactions.

**Results:** The number of unique genes annotated to a chemical ranged from 2 (bromacil) to 5,607 (atrazine), and 121 genes were annotated to SA. In humans, all chemicals tested were highly enriched for SA gene overlap (all p<0.001; parathion PRR=7, cadmium PRR=6.5, lead PRR=3.9, arsenic PRR=3.5, atrazine PRR=2.8). In mice, highest enrichment (p<0.001) was observed for naphthalene (PRR=16.1), cadmium (PRR=12.8), arsenic (PRR=11.6), and carbon tetrachloride (PRR=7.7). In rats, we observed highest enrichment (p<0.001) for cadmium (PRR=8.7), carbon tetrachloride (PRR=8.3), and dieldrin (PRR=5.3). Our findings were robust to 1,000 permutations each of gene sets ranging in size from 100 to 10,000. SA genes were overrepresented in biological processes: inflammatory response (q=0.001), collagen metabolic process (q=1×10^−13^), cell death (q=0.02), and vascular development (q=0.005).

**Conclusion:** We observed chemical gene sets (parathion, cadmium, naphthalene, carbon tetrachloride, arsenic, lead, dieldrin, and atrazine) were highly enriched for SA genes. Exposures to chemicals linked to SA, thus linked to probability of live birth, may deplete fetuses susceptible to adverse birth outcomes. Our findings have critical public health implications for successful pregnancies as well as the interpretation of environmental pregnancy cohort analyses.

## 1. Introduction

Early pregnancy loss through spontaneous abortion (SA), also known as miscarriage, is defined by the loss of a previable fetus prior to 20 weeks of gestation (McNair and Altman 2011). Early pregnancy loss is the most common complication of pregnancy. Estimates of the rate of pregnancies that end in SA typically range from 15 to 20% (Agenor and Bhattacharya 2015; Krieg et al. 2016); however, these values do not include miscarriages or implantation failures that occur before the mother is aware of pregnancy. When these early losses are considered, estimates for early pregnancy loss grow to as high as 50-70% of all pregnancies (Jeve and Davies 2014; Macklon et al. 2002; Salker et al. 2010) and can be an important cause of subfertility or infertility.

Successful pregnancy requires a complex interplay between various immunological, hormonal, and genetic processes (Weselak et al. 2008). A large proportion of miscarriages are explained by fetal chromosomal abnormalities, such as aneuploidy: chromosomal abnormalities affect 50% of first trimester and 20% of second trimester SA fetuses (Choi et al. 2014; Larsen et al. 2013; Tsuiko et al. 2018). Maternal conditions also influence SA risk. Strong casual associations for SA have been identified for acquired maternal thrombophilia and thyroid autoimmunity, as well as other immune or inflammatory related processes (Larsen et al. 2013). Maternal use of substances such as tobacco, cocaine and alcohol during pregnancy have shown significant associations with SA (Ness et al. 1999; Weselak et al. 2008). Environmental toxicant exposures are also associated with SA, including exposure to the pesticide DDT and the heavy metal arsenic (Krieg et al. 2016). Moreover, early pregnancy loss and adverse pregnancy outcomes occur at higher rates in and around contaminated waste sites such as landfills and in areas with drinking water contamination or high air pollution (Grippo et al. 2018; Vrijheid 2000). Relatively little is known however, about the ~80,000 chemicals currently in commerce, and whether they contribute to the risk of SA (Xia et al. 2018). Importantly, exposures are modifiable risk factors and represent an opportunity for SA prevention.

Adverse pregnancy outcomes and reproductive toxicity endpoints are particularly challenging to evaluate in any one *in vitro* or *in vivo* model because they are the result of multiple complex biological processes. In addition, there are key differences between rodents and humans in physiological events involved in pregnancy, such as differences in the production of cytokines and in the process of placentation (Clark 2014; Faas et al. 2005; Moffett and Loke 2006). Women seeking to become pregnant are likely exposed to complex chemical mixtures providing another layer of complexity in hazard identification (Woodruff et al. 2011). Moreover, women may experience pregnancy loss before they are aware of their pregnancy status. In human epidemiologic research, failure to account for a toxicant’s impact on SA can bias potential associations between that toxicant and a birth outcome (Liew et al. 2015). For example, when testing an association between air pollution exposure during pregnancy and autism spectrum disorder diagnosis in children, restricting analyses to live births created a situation with two potential selection-bias processes at play: 1) the chance of live birth is influenced by both air pollution exposure as well as another risk factor for autism; and 2) the depletion of fetuses susceptible to autism from the high exposure group (Raz et al. 2018). Taken together, these factors make evaluating chemical risks to adverse pregnancy outcomes a challenging process. The current analysis is a proof-of-concept study designed to leverage toxicology results in multiple species and prioritize SA-associated chemicals for follow up.

Rigorous interrogation of chemicals and potential pathways involved in SA requires integration of findings from multiple model systems as well as human data. The Comparative Toxicogenomics Database (CTD), developed by North Carolina State University and the National Institute for Environmental Health Sciences (NIEHS), provides a useful tool for such an analysis. The CTD features millions of curated gene-chemical, gene-disease and chemical-disease relationships as well as various other functions (Davis et al. 2017; Davis et al. 2018). To focus our analysis on a set of environmental chemical relevant to a specific geographical location, we selected 25 chemicals found in maternal blood/urine samples, groundwater, tap water, or at Superfund sites in Puerto Rico. Using a combination of CTD data and our own statistical approach, we evaluated the genes associated with each chemical and tested them for enrichment with genes annotated to the CTD disease term “Spontaneous Abortion”. To elucidate potential molecular or cellular mechanisms by which these chemicals could increase risk for SA, we identified Gene Ontology defined Biological Processes associated with specific chemical-gene interactions for a subset of chemicals. The goals of this study were to: 1) identify chemicals associated with increased risk of SA, based on toxicogenomic responses across multiple species (human, mouse and rat) using the CTD; 2) identify chemical impacts on specific molecular targets and cellular pathways involved in SA; and 3) identify targets/pathways commonly impacted by multiple chemicals, suggesting potential toxicant mixtures of concern for this endpoint.

## 2. Methods

### 2.1 Datasets

#### Background gene lists for human, mouse and rat species

The symbols for all genes annotated to each species (*homo sapiens*, *mus musculus* and *rattus norvegicus*) were downloaded from the Comparative Toxicogenomics Database (CTD) (date of download 7/30/18). We then verified that these genes were listed as “active” for each species in the PubMed gene database (queried 8/2/18) (Lu 2011). This returned a total of 42,830 human, 34,321 mouse and 25,150 rat genes.

#### Spontaneous abortion (SA) gene list

We downloaded CTD genes linked to the term “Spontaneous Abortion” (Medical Subject Heading (MeSH) Identifier: D000022, date of download: 7/12/18). This list contains 121 curated human genes. Of these, 111 and 112 have known mouse and rat homologs, respectively. All statistical tests were conducted using human, mouse or rat species data as separate, independent analyses.

#### Chemical gene lists

We selected an optimized number of 25 chemicals for analysis to allow for both 1) analysis of multiple, diverse chemical classes (e.g. metals, organophosphate pesticides, volatile organic compounds, etc.) and 2) feasibility of downstream follow up analysis of chemicals enriched with SA genes. We focused our analysis on chemicals relevant to exposures in the population of greatest concern for the SA endpoint (i.e., pregnant women) within a specific geographical location (Puerto Rico), a region with high rates of adverse pregnancy outcomes (Martin et al. 2018) and extensive contamination with Superfund chemicals (USEPA, 2018). Therefore, we selected chemicals based on their presence in maternal blood/urine samples, groundwater, tap water, Superfund sites or reported usage in Puerto Rico. The total list of 25 chemicals included heavy metals [arsenic, cadmium, lead and chromium (Padilla et al. 2011)]; phthalate esters [dibutyl phthalate, diethyl phthalate and diethylhexyl phthalate (Cantonwine et al. 2014; Padilla et al. 2011)], volatile organic compounds [perchloroethylene, trichloroethylene, carbon tetrachloride, chloroform and methylene chloride (Padilla et al. 2011)], pesticides [aldrin, atrazine, bromacil, dieldrin, glyphosate, methyl parathion and parathion (Irizarry et al. 1989; Marnio et al. 2016; Padilla et al. 2011)], phenols [triclosan and benzophenone-3 (Meeker et al. 2013)], polycyclic aromatic hydrocarbons [naphthalene, phenanthrene and pyrene (Cathey et al. 2018)] and a personal care product [N,N-Diethyl-meta-toluamide (Lewis et al. 2014)]. Gene lists for these chemicals were downloaded for each of three species (human, mouse and rat) from the CTD website on 9/29/18. Bromacil was dropped for further analysis due to a having an exceptionally small gene list (n=2 genes).

### 2.2 Statistical tests

#### Enrichment testing of chemical gene lists and SA genes

All analyses were conducted in R statistical software. Code used to conduct analyses and generate figures and tables is publicly available (https://github.com/bakulskilab). We conducted univariate descriptive statistics on each species-specific chemical gene list. Within species, we generated 2×2 descriptive tables for genes associated with each chemical by genes associated with SA. Based on the gene distributions for each chemical and each species, we selected appropriate enrichment test statistics. When expected frequencies per cell were greater than five, enrichment testing was conducted using standard chi-square test for independence. When the expected frequency of genes per cell was less than five and greater than one, we used the ‘N-1’ chi-squared test (Campbell 2007). Chemicals with expected frequency less than one in any cell of the 2×2 table were dropped from further analysis for that species. Out of the 24 remaining chemicals × three species (72 possible comparisons), we proceeded with 20 enrichment tests. See **Supplementary Figure 1** for a flow chart illustrating chemical inclusions/exclusions. We considered p<0.05 to be suggestively enriched and a Bonferroni correction was used to account for multiple comparisons (n=20 tests; p<0.0025).

#### Sensitivity analyses

We used Fisher’s exact tests to confirm chemical-disease gene enrichment associations observed in the primary analyses. The gene sample size was always greater than 1,000 in this study, thus we prioritized the chi-squared test and used Fisher exact test as a comparison. For all chemicals showing significant gene-disease/gene-chemical associations, the proportional reporting ratio (PRR) was calculated to identify the direction and magnitude of enrichment (i.e. more overlapping genes than expected by chance) or depletion (i.e. fewer overlapping genes than expected by chance). A PRR>1 indicates enrichment while a PRR<1 indicates depletion.

#### Cross-validation of enrichment results

To assess the susceptibility of the SA gene list to false-positives, we performed cross-validation for enrichment testing with SA genes. We randomly permuted 1000 pseudo-chemical gene lists each of specific sizes (100, 500, 1000, 2500, 5000 and 10000 genes) to reflect the range of genes associated with chemicals in our dataset. Genes in these random lists were then tested for enrichment with the exact SA genes. We calculated then number of permuted tests meeting our significance criteria by chance. We visualized the patterns in results with a density plot of observed association p-values.

#### Gene Ontology (GO) enrichment analysis of SA genes

To gain insight into specific biological functions represented in the 121 genes contained in the CTD term “Spontaneous Abortion” (SA), we identified enriched Gene Ontology terms represented by the 121 SA genes using the DAVID (Database for Annotation, Visualization, and Integrated Discovery) pathway enrichment tool (Huang da et al. 2009a, b). The 121 gene symbols were used as input for the enrichment and “homo sapiens” was used as the background or reference gene list. We filtered SA gene enriched Gene Ontology terms with the REVIGO online analysis tool. REVIGO provides a more easily interpretable Gene Ontology list by removing redundant Gene Ontology terms (Supek et al. 2011).

#### Identification of chemical-gene associations for enriched Gene Ontology terms

Based on enrichment testing results, we prioritized a subset of five chemicals that were enriched from the human chemical data (arsenic, cadmium, lead, atrazine and parathion). We visualized overlap in SA genes interacting with these chemicals using a heatmap. Gene membership in each of four selected Gene Ontology terms was highlighted.

## 3. Results

### 3.1 Gene-chemical associations

We surveyed 25 chemicals for their associated genes in the CTD and noted that the number of unique genes per chemical ranged from two (bromacil) to 5,607 (atrazine). We observed a mean of 1,441 genes per chemical and a median of 784 genes per chemical (**Supplementary Table 1**). The proportion of genes associated with each chemical varied by species (human, rat, mouse) (Figure 1). Parathion had the highest proportion of genes (95%) from obtained human studies and chloroform had the lowest proportion (13%). Twenty-three out of the 25 chemicals analyzed included genomic data from all three species. Benzophenone-3 included data from human and rat. Bromacil included data from human only.

**Figure 1.**
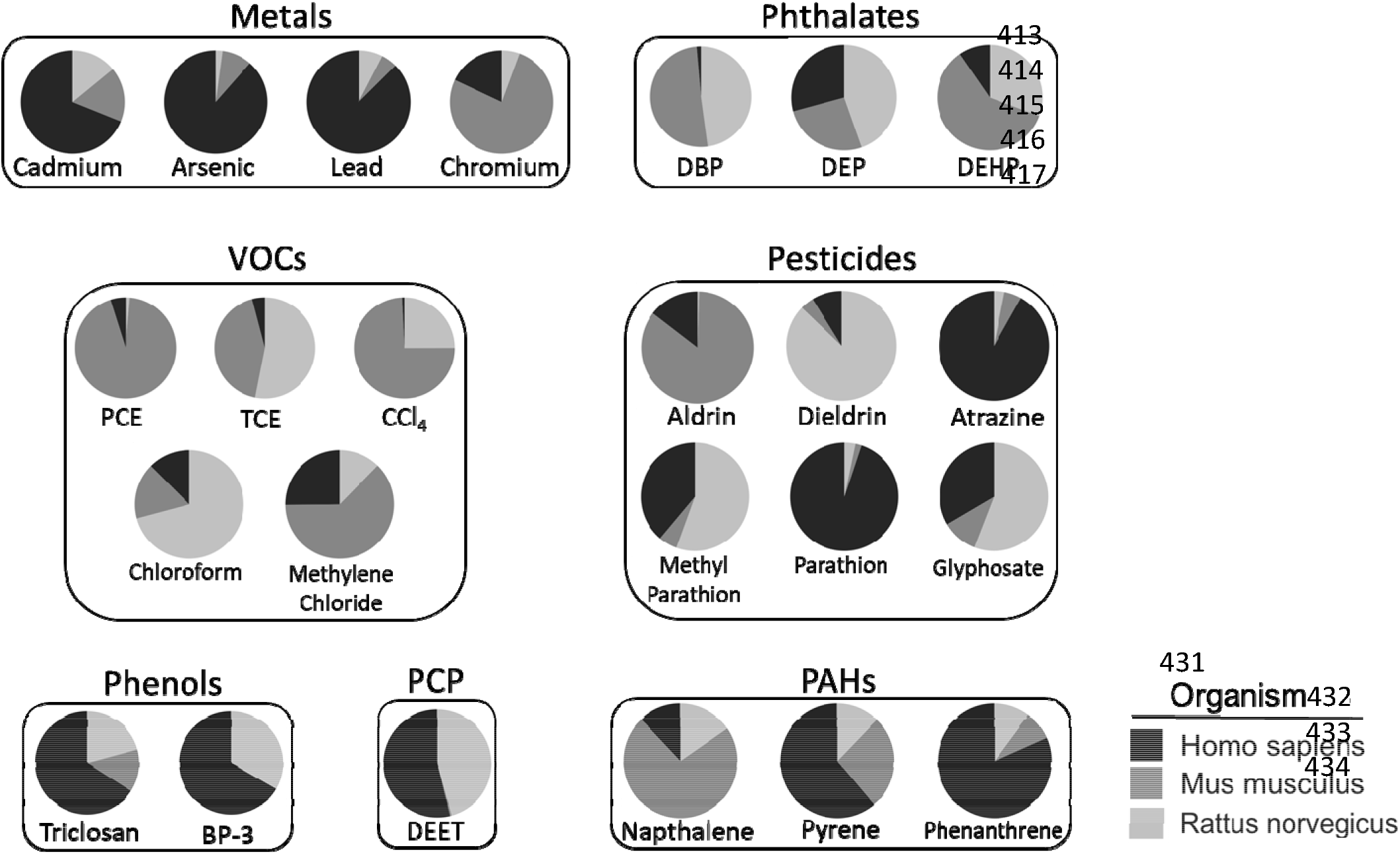
Proportion of genes obtained for each chemical by species. Lists of genes found to interact with 24 different chemicals in any of 3 different species (homo sapiens, mus musculus and rattus norvegicus) were downloaded from the Comparative Toxicogenomics Database (date of download: 7/30/18). The proportion of genes associated with each species varied across compounds with parathion having the highest (95%) and chloroform the lowest (13%) proportion of genes obtained from homo sapiens. VOCs: volatile organic carbons, PCE: perchloroethylene, TCE: trichloroethylene, CCl_4_: carbon tetrachloride, BP-3: benzophenone-3, DEET: N,N-Diethyl-meta-toluamide, PCP: personal care product, PAHs: polycyclic aromatic hydrocarbons

We filtered chemicals by species based on the number of associated genes. Bromacil was removed from consideration for all species. More chemicals were removed due to an expected value for Chi-squared testing of <1 (19, 15 and 18 chemicals for human, mouse and rat species, respectively). This left 13 total chemicals that were tested for enrichment with SA genes in at least one species. These chemicals were: cadmium, arsenic, lead, chromium, DBP, DEHP, PCE, TCE, CCl_4_, dieldrin, atrazine, parathion and naphthalene.

### 3.2 Enrichment testing for chemical associations with SA

Twelve of the 13 chemicals tested were enriched with SA genes (corrected chi-squared p<0.05) in one or more species. Perchloroethylene was tested in the mouse data but not significantly enriched (**Table 1**). Cadmium was the only chemical enriched across all three species. Of the compounds tested, five chemicals were enriched for human, seven for mouse and six for rat. Metals had the highest percentage of chemicals with 4/4 (100%) enriched for SA genes in at least one species. This was followed by phthalates with 2/3 (66.7%) enriched compounds, then pesticides (3/6, 50%), VOCs (2/5, 40%), PAHs (1/3, 33%), phenols (0/2, 0%) and the personal pesticide product DEET (0/1, 0%). The PPRs>1 for these associations reflect higher risk of SA due to more overlap with SA genes (as opposed to depletion or lower risk) (Figure 2). The highest PRRs were observed for arsenic (11.6), cadmium (12.8) and naphthalene (16.1). The confidence intervals for some chemicals were relatively large (e.g. cadmium, dieldrin) (**Supplementary Table 2**) suggesting a higher degree uncertainty around the magnitude of enrichment for these compounds.

**Figure 2.**
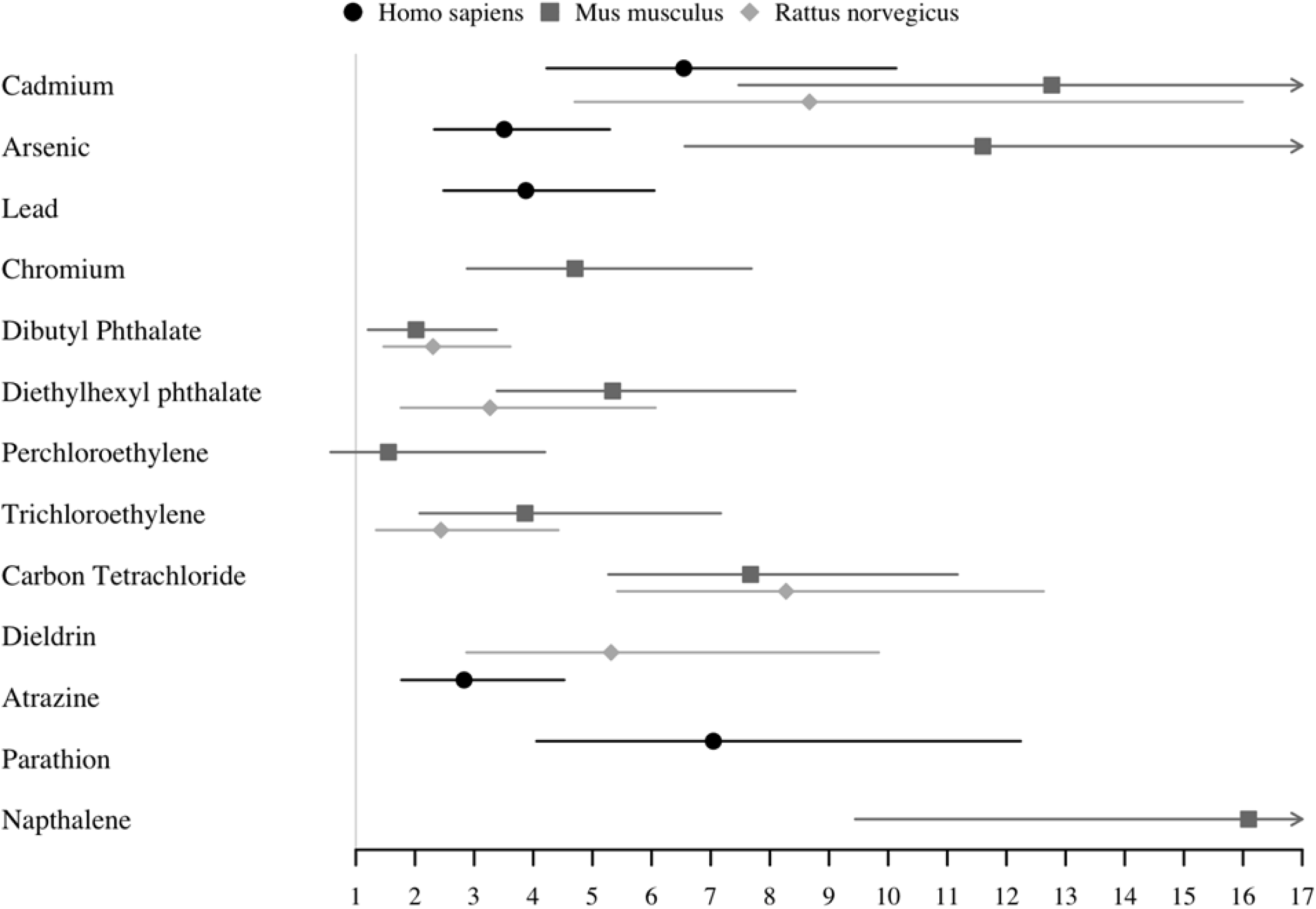
Proportional reporting ratios (PRRs) and confidence intervals for chemical associations with spontaneous abortion (SA) genes. Proportional reporting ratios were calculated for chemicals significantly enriched with SA genes in order to identify the direction and magnitude of enrichment or depletion. Errors show 95% confidence intervals.

### 3.3 Cross-validation

For randomly permuted human gene data, 0.37% of tests achieved chi-square corrected p-value<0.05 across all gene set sizes (n=100, 500, 100, 2500, or 5000) and 0.34% achieved Fisher’s test corrected p-value<0.05. For randomly permuted mouse gene data, 0.39% of tests achieved chi-square corrected p-value<0.05 and 0.40% achieved Fisher’s test corrected p-value<0.05. For randomly permuted rat gene data, 0.40% of tests achieved chi-square corrected p<0.05 and 0.35% of tests achieved Fisher’s test corrected p-value<0.05. Cross-validation results across all p-value thresholds for human data are shown in **Supplementary Figure 2**.

### 3.4 Functional analysis of Spontaneous Abortion gene list

Among the SA genes, pathway enrichment analysis identified 143 enriched Gene Ontology terms (FDR<0.05). After removal of redundant terms using REVIGO, 46 enriched terms remained. Based on these results, we identified the following enriched Gene Ontology terms based known links to processes involved in SA: “collagen metabolic process” (GO:0032963), “inflammatory response” (GO:0006954), “cell death” (GO:0008219) and “vasculature development” (GO:0001944). Consistent with the known etiology of SA, enriched terms included processes associated with inflammation and immune processes (“inflammatory response”, GO:0006954, p=0.001; “immune response”, GO:0006955, p=0.0001), collagen metabolism (“collagen metabolic process”, GO:0032963, p=1×10^−13^), cell death (“cell death”, GO:0008219, p=0.02) and tissue development (“vasculature development”, GO:0001944, p=0.0005; “skeletal system development”, GO:0001501, p=0.02).

### 3.5 Overlap of genes targeted by enriched chemicals and functional analysis

We prioritized the five chemicals enriched in the human data for further study: arsenic, cadmium, lead, atrazine and parathion. Each of the five chemicals are shown to impact multiple genes in the SA gene list (Figure 3). Some genes are impacted by multiple chemicals, such as *CYP1A1* and *NCAM1* that were impacted by all five chemicals. Other genes are unique to specific chemicals, such as *IL24* and *JAK2* (specific to arsenic) or *COL6A1* and *LAMA4* (specific to atrazine). We observed chemical-gene interactions across all four of the Gene Ontology terms investigated. Genes did not appear to cluster by any particular Gene Ontology process or chemical. *CYP1A1* and *NCAM1* were not involved in the Gene Ontology processes investigated.

**Figure 3.**
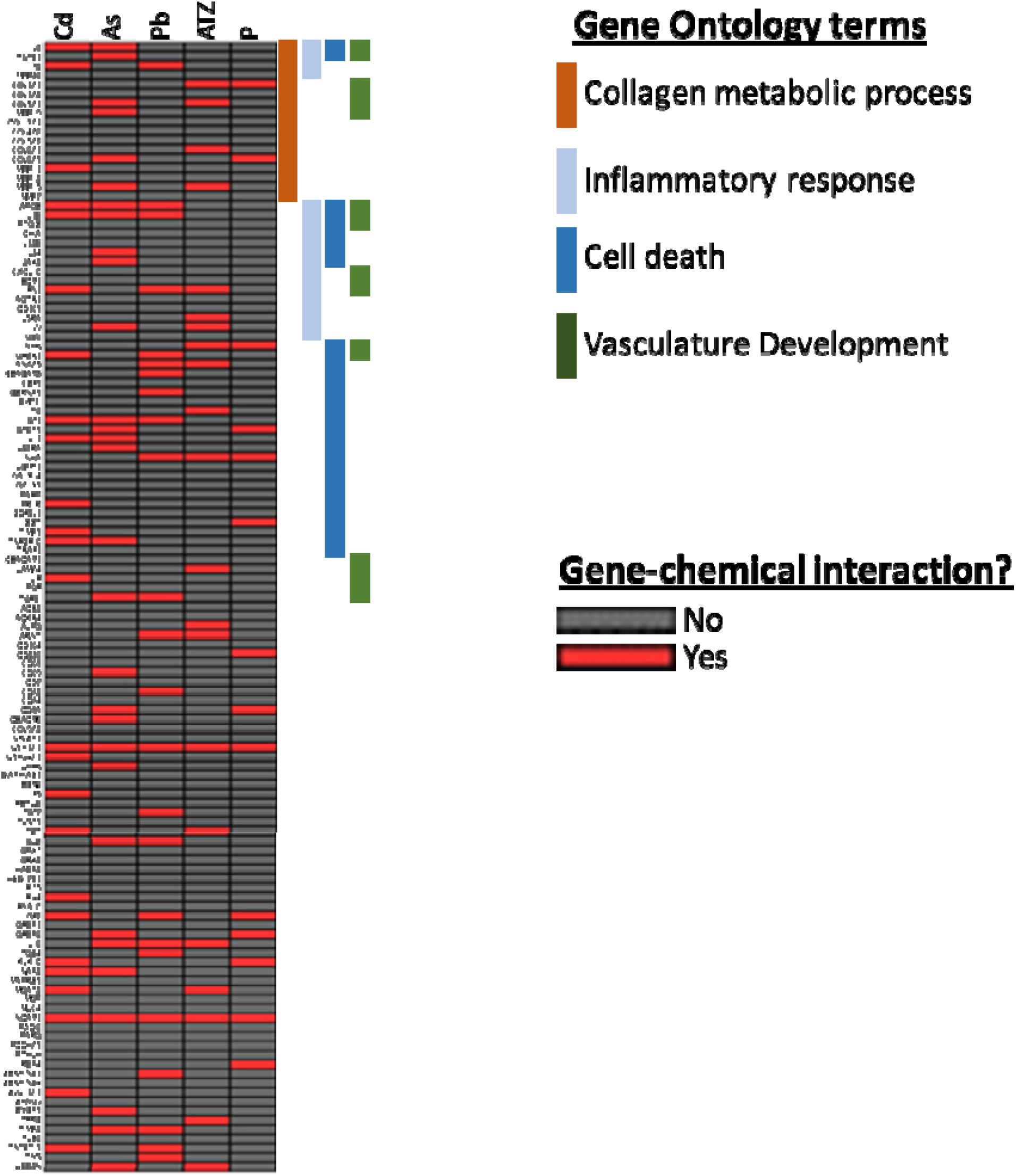
Chemical-gene interactions for 121 Spontaneous Abortion (SA) genes and associations with selected Gene Ontology terms. Chemical interactions for 121 SA genes were plotted for 5 chemicals (Cd: cadmium, As: arsenic, Pb: lead, ATZ: atrazine and P: parathion). Associations with select Gene Ontology terms relevant to the etiology of SA (collagen metabolic process, inflammatory response, cell death and vasculature development) were plotted in parallel in order to identify potential toxicological mechanisms linking chemical exposure to SA.

## 4. Discussion

The present study evaluated a set of chemicals detected in Superfund sites and in pregnant women for associations with molecular markers of spontaneous abortion using publicly available toxicogenomic data contained in the CTD. Our results showed that a diverse set of chemicals representing metals, pesticides, volatile organic carbons and polycyclic aromatic hydrocarbons affect more SA genes than would be expected by chance in at least one of the species studied (human, mouse or rat).

Our results are consistent with epidemiological literature showing associations between heavy metal exposure and SA. For example, in our study arsenic, cadmium, chromium and lead all were all highly enriched with SA genes. Consistent with these results, multiple meta-analyses of epidemiological literature have concluded that high levels of arsenic exposure (>50 ppb) are associated with SA and that plausible mechanisms exist for arsenic causing SA due to its endocrine disrupting properties (Milton et al. 2017; Rahman et al. 2016). A recent report showed lead levels in the hair of pregnant woman were associated with risk of missed abortion (i.e., loss of pregnancy with retention of products of conception) (Zhao et al. 2017). Similarly, increased incidence of SA has also been reported in women with occupational exposure to chromium (Remy et al. 2017). Animal studies have found that cadmium decreases incidence of pregnancy through decreased implantation and fetal resorbtions (Jacobo-Estrada et al. 2017). Cadmium impacts on pregnancy outcomes are plausible because cadmium is an endocrine disruptor which is known to decrease levels of hormones necessary to maintain pregnancy, such as progesterone (Jacobo-Estrada et al. 2017; Nampoothiri and Gupta 2008; Piasek and Laskey 1994). However, Buck-Louis et al. reported no association with pregnancy loss at environmentally relevant levels of exposure to cadmium in humans (Buck Louis et al. 2017). The results of our study contribute to the body of evidence that exposure to arsenic, cadmium, chromium and lead constitute a risk factor for SA, although the threshold of exposure leading to increased risk is not well characterized.

Two of the phthalates we tested were also enriched for SA genes, with DBP enriched for rat and DEHP enriched for mouse. These results are consistent with several reports in human observational studies. Both DBP and DEHP levels in maternal hair samples were associated with missed abortion (Zhao et al. 2017). In addition, urinary levels of the main metabolite of DEHP (monoethylhexyl phthalate) were higher in women with missed abortion (Yi et al. 2016) and in women with pregnancy loss undergoing medically assisted reproduction (Messerlian et al. 2016; Toft et al. 2012; Zhao et al. 2017). In another study, a metabolite of DBP (monobutyl phthalate) was not associated with recurrent SA, although a metabolite of a structurally similar phthalate (mono-isobutyl phthalate) was associated with recurrent SA (Peng et al. 2016). Phthalatess are well known endocrine disruptors (Qureshi et al. 2016). Phthalates also generate oxidative stress (Tetz et al. 2013; Wang et al. 2010), inflammation (Wang et al. 2010) and epigenetic changes (Grindler et al. 2018; Strakovsky and Schantz 2018) in gestational tissues and cells in culture, suggesting several potential mechanisms to increase risk of SA. Clarifying these mechanisms is worthy of further study.

Consistent with our results, carbon tetrachloride has been shown to induce abortion in a rat model (Narotsky et al. 1997). While not known to cause SA specifically, TCE has been shown to target female germ cells by decreasing fertilizability of rat oocytes (Narotsky et al. 1997; Wu and Berger 2007). Although there is less epidemiological evidence for associations with SA in humans, both compounds are associated with low birth weight and birth defects (Bove et al. 1995; Forand et al. 2012; Ruckart et al. 2014). Similarly, while epidemiological and experimental data link the pesticides atrazine, dieldrin and parathion (enriched for SA in our study) to certain adverse pregnancy outcomes, evidence for SA is mixed. For example, the spouses of workers who used higher levels of atrazine were at increased risk of SA (Petrelli et al. 2003), though a recent systematic review of the literature found a cumulative lack of evidence linking atrazine to adverse birth outcomes, including SA (Goodman et al. 2014). Similarly, whereas dieldrin is associated with adverse outcomes in pregnancy such as gestational hypertension (Savitz et al. 2014) and altered thyroid hormone levels in cord blood (Luo et al. 2017), another study found no association between maternal serum levels and missed abortion (Bercovici et al. 1983). Parathion does not affect the number of live fetuses/litter in rats (Keith et al. 2017), but one small study in pregnant women (n=20) showed altered placental morphology in an exposed group, suggesting impacts on gestational tissues that may have implications for SA (Levario-Carrillo et al. 2001). Thus, the VOC and pesticide enrichment for genomic markers of SA may reflect responses in gestational tissues that are not linked to SA specifically but may have implications for other pregnancy outcomes. Further studies may shed light on the effects on these chemicals and determine if exposures have any implications for risk of SA.

SA is a multi-factor process involving a variety of signaling pathways. We conducted an analysis of enriched Gene Ontology terms for the SA genes in the CTD in order to determine which specific pathways were overrepresented among these 121 genes. Interestingly, our results were consistent with known etiologies of SA and in some cases were reflective of the modes of toxicity for the chemicals studied. For example. we observed that the Gene Ontology term “collagen metabolic process” was enriched among SA genes, consistent with research showing women with a history of spontaneous abortions had decreased levels of collagen types IV and V in the decidua (Iwahashi et al. 1996; Iwahashi and Nakano 1998). Notably, several of the metals significantly associated with SA also disrupt collagen metabolism in animal models, offering insight into potential modes of action by which these compounds could present a hazard for this endpoint. Arsenic exposure (50 ppb in drinking water) decreases collagen gene expression and disrupts collagen deposition in the lungs and heart of mice (Hays et al. 2008). Cadmium disrupts collagen metabolism in the bones of rats (Galicka et al. 2004) and lead disrupts collagen metabolism in the brain capillaries in calves (Ahrens 1993). Less is known about the effects of the pesticides atrazine and parathion on collagen metabolism, but a study of male reproductive effects of atrazine exposure in rats noted reduced testicular collagen fiber (Kniewald et al. 2000). Whether exposure to these the chemicals leads to increased risk for SA via disrupted collagen metabolism in the decidua warrants further study.

SA genes were also enriched for the Gene Ontology term “inflammatory response”. The role of inflammation in implantation, pregnancy and parturition is complex with both pro and anti-inflammatory cytokines playing various roles. However, an overall shift from Th2 (anti-inflammatory) to Th1 (pro-inflammatory) response in the decidua is a potential cause for SA (Berger 2000; Romero et al. 2004) and increased levels of cytokines like interferon-γ (IFNγ), IL-2 and TNFα are measured in women with recurrent SA (Ng et al. 2002; Pandey et al. 2005). Epidemiology studies have shown arsenic (Ahmed et al. 2011; Dutta et al. 2015; Prasad and Sinha 2017), cadmium (Everson et al. 2018; Lin et al. 2009; Messner and Bernhard 2010) and lead (Machon-Grecka et al. 2018; Sirivarasai et al. 2013; Wang et al. 2009) are associated with increased markers of inflammation, including in gestational tissues such as the placenta. For example, cadmium increases blood levels of complement component 5 fragment (C5a), which is associated with miscarriage (Jacobo-Estrada et al. 2017; Zhang et al. 2016). In comparison, although parathion is linked to asthma (characterized by airway inflammation) (Hoppin et al. 2009) and atrazine causes inflammation in the prostate of rats (Scialli et al. 2014), to our knowledge no studies have reported these compounds causing inflammation in gestational tissues. Future experiments may clarify if this is a relevant mechanism of toxicity for these compounds in regard to SA.

“Cell death” was another Gene Ontology term enriched among SA genes. Apoptosis and cell death is associated with SA. Placentas from women with recurrent SA have increased levels of apoptosis compared to women with normal pregnancies (Atia 2017). Exposures to toxic chemicals such as polycyclic aromatic hydrocarbons induce apoptotic pathways and lead to embryonic loss in animal models, a proposed mechanism explaining higher miscarriage rates in women who smoke (Detmar et al. 2006). Arsenic and atrazine cause apoptosis in mouse embryo models (Liu et al. 2003; Scialli et al. 2014), and lead and cadmium cause apoptosis in the placenta of rats (Erboga and Kanter 2016; Wang et al. 2014), consistent with our results showing that these compounds impact multiple SA genes involved in cell death.

Finally, the gene lists for all five of the chemicals enriched for SA in the human data contain genes in the Gene Ontology term “vasculature development”. Vasculature development at the fetal-maternal interface is critical to the maintenance of pregnancy. Women undergoing SA have decreases in vascularization in chorionic villi (Meegdes et al. 1988). Arsenic exposure disrupts this process in mouse models, leading to SA (He et al. 2007). Cadmium disrupts vasculature development in zebrafish (Cheng et al. 2001) and in disrupts growth and migration in human vascular endothelial cells (Kishimoto et al. 1996). Lead has similar effects on the rat blood-brain barrier (Wang et al. 2007). While atrazine or parathion both interact with multiple genes in the vasculature development Gene Ontology term, to our knowledge there is currently little reported evidence for effects of on the vasculature developmental process in gestational tissues.

Contrary to our initial hypothesis, genes did group either by chemical class (i.e., metals vs. pesticides) or by Gene Ontology term. Instead, all five analyzed chemicals impacted genes across one or more of the Gene Ontology terms studied. Taken together our results highlight that diverse chemicals can affect multiple molecular targets within biological pathways relevant to mechanisms underlying SA, and that publicly available databases can be used to “scan” a wide range of chemicals for potential interactions with these targets. Given the biological complexity of reproductive and developmental disorders and the inherent difficulties in replicating them in animal and cell models, the methods used in this study could be used for hazard identification for toxicant impacts on reproductive/developmental endpoints. In addition, this approach could prove especially useful for assessing complex mixtures of chemicals such as those found at Superfund or fracking wastewater sites by identifying a set of chemicals in a particular mixture that are likely to affect the same set of pathways underpinning a known toxicological endpoint or disease.

Spontaneous abortion (SA) is a notable toxicological endpoint for reproductive epidemiology studies because toxicant exposure that leads to increased rates of SA may be biased toward the null when investigating other health effects. Weisskopf et al. demonstrated that biases toward the null hypothesis of no effect can be introduced through the “depletion of susceptibles” in environmental epidemiology studies. That is, individuals that participate in exposure-health studies may be a select group of individuals who are resistant to toxicant induced health effects whereas those more sensitive may not be included (Weisskopf et al. 2015). This situation could arise in the case of chemicals contributing to SA, because sensitive pregnancies may be lost before subject recruitment into epidemiology studies. Quantitative bias analysis may be needed to assess the potential influence of live birth bias in birth outcome or developmental studies (Lash et al. 2014; Weuve et al. 2018). Our results suggest that several chemicals that we investigated could generate such biases.

Strengths of this study and our statistical approach include the ability to screen a large number of chemicals for potential risks to a highly complex toxicological endpoint in a relatively short amount of time. Moreover, by analyzing the Gene Ontology terms represented among SA genes we were able to gain insight into specific mechanisms by which chemicals might contribute to increased risk of SA, providing more biological coherence to our interpretations than would be achieved by simply testing chemicals for associations with disease terms. In addition, before conducting our enrichment tests, we performed cross-validation in order to verify that the SA gene list was resistant to false-positives. Finally, by conducting enrichment testing in each of the three species (human, mouse and rat) separately, we were able to focus our downstream analysis on chemicals that were most relevant to currently available human data (i.e. cadmium, arsenic, lead, atrazine and parathion) and follow up with chemicals enriched in other species in a later analysis. In addition, this allowed us to identify one compound (cadmium) as being enriched for SA across all three species we tested (human, mouse and rat), illustrating a potentially conserved response across several mammalian species.

There are also several limitations to the current study that should be noted. The results presented demonstrate statistical associations between genes impacted by chemicals and those involved in SA. This analysis was not able to address matters of dose-response or directionality of gene expression changes (up- or down-regulation) for these chemicals. Therefore, this study is meant as a demonstration of a method of hazard identification and a not dose-response study. The lack of tissue specificity is another limitation that should be noted, as the chemical-gene lists were obtained from a wide variety of studies using various tissues which may not reflect the chemical-gene interactions occurring in female reproductive tissues relevant to the physiological processes of SA. Future studies will address these limitations, through use of cell models or additional databases.

Taken together, our results contribute to the body of evidence that exposure to several classes of chemicals, including metals and pesticides, constitute risk factors for SA. Our results show that publicly available databases, such as the CTD, can be used to screen a large number of chemicals for effects on molecular responses related to a complex reproductive endpoint. This approach identified multiple compounds that affect more SA genes than would be expected by chance and provided insight into which normal cellular pathways are targeted by these toxicants. This method provides a basis for considering mixture effects (synergistic, additive, etc.) when assessing the risks these chemicals pose for SA by allowing us to identify toxicants that target similar diseases and cellular pathways, thus providing key insight into disease-chemical relationships and common modes of action across chemicals. Our findings have critical implications for the design and interpretation of pregnancy cohort analyses with environmental exposures because chemicals contributing to SA could lead to sensitive pregnancies being lost before recruitment into epidemiology studies.

## Funding acknowledgements

This research was supported by the Michigan Center on Lifestage Environmental Exposures and Disease (P30 ES017885). Drs. Harris, Meeker, and Loch-Caruso were supported by grants from the National Institutes of Health, Superfund Research Program (P42 ES017198). Additional funding for Dr. Harris was provided by the National Center for Advancing Translational Sciences (UL1TR002240). Dr. Bakulski was supported by grants from the National Institutes of Health (R01 ES025531; R01 ES025574; R01 AG055406; R01 MD013299).

## Notes

Competing Financial Interests: The authors declare they have no actual or potential competing financial interests.

https://ctdbase.org/

