## Supplementary Material for "Toxicants Associated with Spontaneous Abortion in the Comparative Toxicogenomics Database (CTD)"

2 Supplementary Tables

2 Supplementary Figures

**Supplementary Table 1.** Number of unique genes by species associated with 25 chemicals in the Comparative Toxicogenomics Database.

|  | **# of unique genes** | | | |
| --- | --- | --- | --- | --- |
| **Chemical** | **All Species** | **Human** | **Rat** | **Mouse** |
| Cadmium | 2,549 | 1,682 | 343 | 416 |
| Arsenic | 4,034 | 3,679 | 89 | 388 |
| Lead | 2,965 | 2,604 | 225 | 148 |
| Chromium | 1,790 | 344 | 107 | 1,461 |
| DBP | 4,784 | 77 | 2,674 | 2,839 |
| DEP | 59 | 19 | 29 | 17 |
| DEHP | 2,459 | 271 | 865 | 1,686 |
| PCE | 853 | 42 | 12 | 808 |
| TCE | 2,167 | 91 | 1,180 | 951 |
| CCl4 | 3,520 | 29 | 974 | 2,898 |
| Chloroform | 93 | 12 | 68 | 16 |
| Methylene chloride | 191 | 49 | 24 | 122 |
| Triclosan | 746 | 89 | 28 | 18 |
| BP3 | 84 | 6 | 3 | 0 |
| Aldrin | 224 | 32 | 1 | 187 |
| Dieldrin | 1,368 | 51 | 505 | 23 |
| Atrazine | 5,607 | 2,961 | 100 | 163 |
| Methyl parathion | 90 | 21 | 30 | 3 |
| Parathion | 822 | 781 | 27 | 15 |
| Glyphosate | 556 | 88 | 147 | 28 |
| Napthalene | 454 | 55 | 70 | 330 |
| Pyrene | 277 | 30 | 6 | 13 |
| Phenanthrene | 130 | 40 | 5 | 4 |
| DEET | 203 | 114 | 97 | 1 |
| Bromacil | 2 | 2 | 0 | 0 |
| **Mean** | **1,441** | **526.8** | **304.4** | **501.4** |
| **Median** | **746** | **55** | **70** | **122** |

**Supplementary Table 2.** Results from tests for enrichment of chemical gene lines with spontaneous abortion genes for three species.

|  | **Chemical** | **Enrichment p-value** | **PRR (CI)** |
| --- | --- | --- | --- |
| Human | Cadmium | <0.001^a^ | 6.5 (4.2-10.1) |
|  | Arsenic | <0.001^b^ | 3.5 (2.3-5.3) |
|  | Lead | <0.001^b^ | 3.9 (2.5-6.0) |
|  | Atrazine | <0.001^b^ | 2.8 (1.8-4.5) |
|  | Parathion | <0.001^a^ | 7.0 (4.1-12.2) |
| Mouse | Cadmium | <0.001^a^ | 12.8 (7.5-21.8) |
|  | Arsenic | <0.001^a^ | 11.6 (6.6-20.5) |
|  | Chromium | <0.001^a^ | 4.7 (2.9-7.7) |
|  | DBP | 0.058^b^ | 2.1 (1.2-3.4) |
|  | DEHP | <0.001^b^ | 5.3 (3.4-8.4) |
|  | PCE | 1^a^ | 1.55 (0.6-4.2) |
|  | TCE | <0.001^a^ | 3.9 (2.1-7.2) |
|  | CCl_4_ | <0.001^b^ | 7.7 (5.3-11.2) |
|  | Napthalene | <0.001^a^ | 16.1 (9.4-27.4) |
| Rat | Cadmium | <0.001^a^ | 8.7 (4.7-16.0) |
|  | DBP | 0.001^b^ | 2.3 (1.5-3.6) |
|  | DEHP | 0.0004^a^ | 3.2 (1.8-6.1) |
|  | TCE | 0.015^b^ | 2.4 (1.3-4.4) |
|  | CCl_4_ | <0.001^a^ | 8.3 (5.4-12.6) |
|  | Dieldrin  ^a^“N-1” chi-squared test  ^b^Standard chi-squared test  CI: confidence interval  PRR: Proportional reporting ratio | <0.001^a^ | 5.3 (2.9-9.8) |

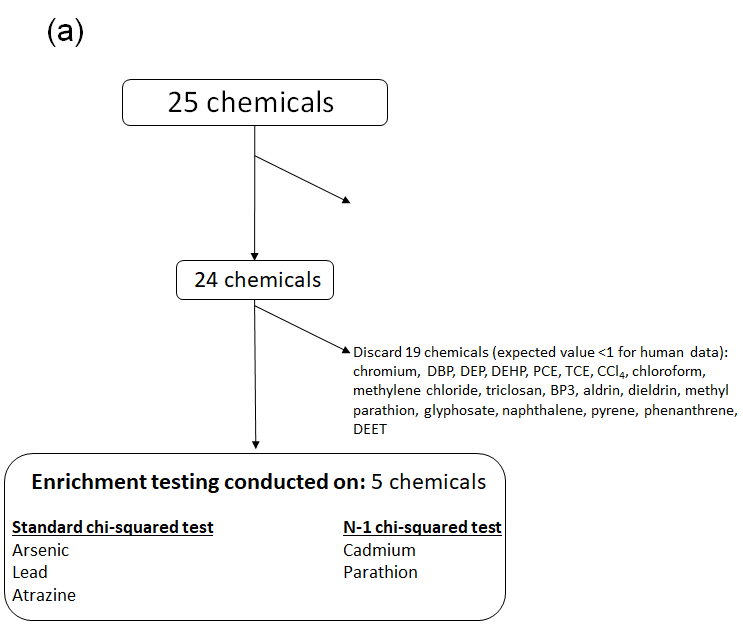

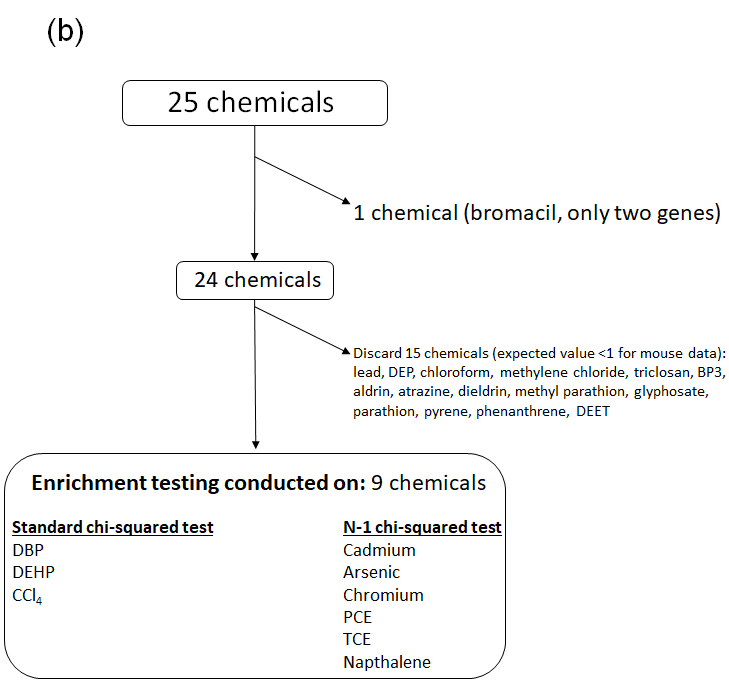

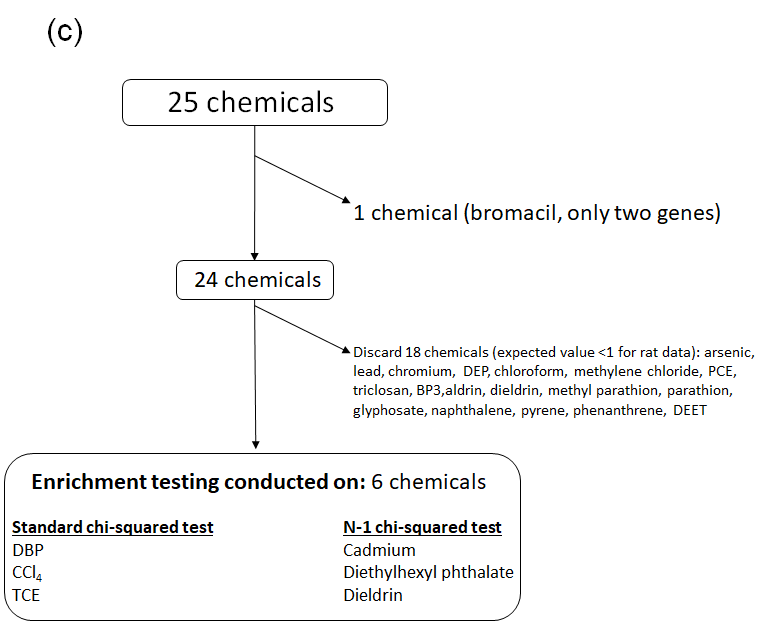

**Supplementary Figure 1.** Flow chart for chemical inclusion/exclusion for enrichment testing for **(a)** human, **(b)** mouse and **(c)** rat species. BP3: benzophenone-3; CCl_4_: carbon tetrachloride, DEET: N,N-Diethyl-meta-toluamide, DBP: dibutyl phthalate, DEP: diethyl phthalate, DEHP: diethylhexyl phthalate, PCE: perchloroethylene, TCE: trichloroethylene

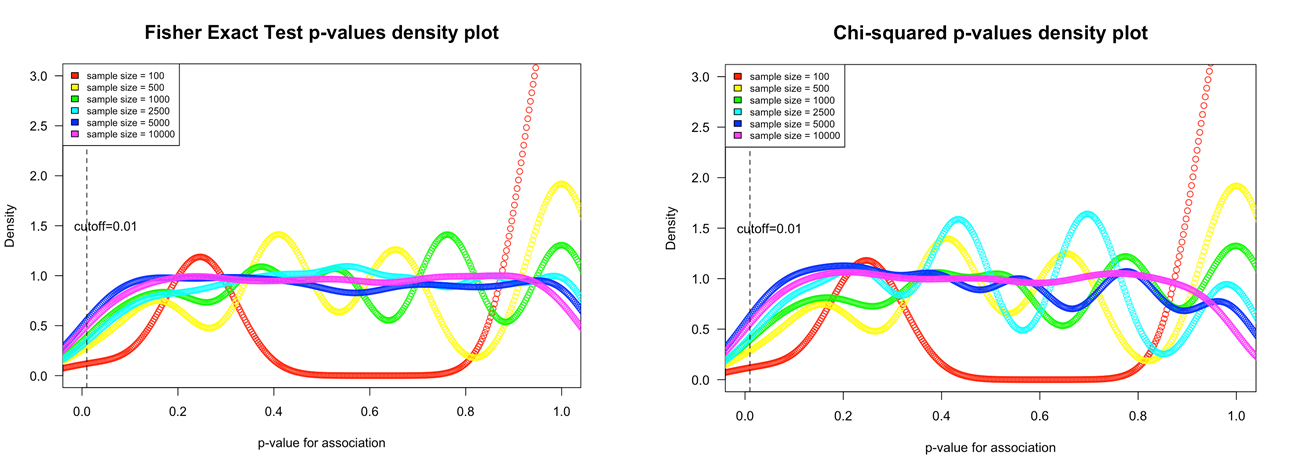

(b)

(a)

**Supplementary Figure 2. Cross-validation of spontaneous abortion findings.** To test the robustness of the spontaneous abortion findings against false positives, we performed cross-validation for enrichment using simulated chemical gene lists, randomly selected and of various list sizes (100, 500, 1000, 2500, 5000 and 10000 genes). We permuted the simulated chemical gene selection 1,000 times per gene list size. We tested for enrichment of the spontaneous abortion gene list with the simulated chemical gene lists using both Fisher exact (a) and Chi-squared tests (b).
